# Automated design of CRISPR prime editors for thousands of human pathogenic variants

**DOI:** 10.1101/2020.05.07.083444

**Authors:** John A. Morris, Jahan A. Rahman, Xinyi Guo, Neville E. Sanjana

## Abstract

Prime editors (PEs) are CRISPR-based genome engineering tools that can introduce precise base-pair edits at specific locations in the genome. These programmable gene editors have been predicted to repair 89% of known human pathogenic variants in the ClinVar database, although these PE constructs do not presently exist. Towards this end, we developed an automated pipeline to correct (therapeutic editing) or introduce (disease modeling) human pathogenic variants that optimizes the design of several RNA constructs required for prime editing and avoids predicted off-targets in the human genome. However, using optimal PE design criteria, we find that only a small fraction of these pathogenic variants can be targeted. Through the use of alternative Cas9 enzymes and extended templates, we increase the number of targetable pathogenic variants to >50,000 variants and make these pre-designed PE constructs accessible through a web-based portal (http://primeedit.nygenome.org). Given the tremendous potential for therapeutic gene editing, we also assessed the possibility of developing universal PE constructs. By examining the overlap of different PE components with common human genetic variants in dbSNP, we find that common variants affect only a small minority of designed PEs.

---

Recently, Anzalone *et al*.^1^ developed prime editors (PEs) for introducing precise edits with base-pair resolution using a Cas9 nickase tethered to a reverse transcriptase. PEs are targeted via a PE guide RNA (pegRNA) to a specific genomic locus, where they make a targeted single-stranded break (nick) in the DNA and reverse transcribe a repair template on the 3’ end of the pegRNA with the desired base-pair substitution, insertion or deletion. Through an analysis of the ClinVar database of human genetic variants^2^, it was suggested that prime editing may be able to correct up to 89% of known genetic variants associated with human diseases^1^. Here, we developed a computational pipeline to design prime editors to correct pathogenic variants in ClinVar for therapeutic gene editing and to introduce these variants into wild-type cells to create disease models (**Fig. 1**). We then present PE design considerations to further improve the number of targetable variants and prime editor reagents per variant.

**Figure 1.**
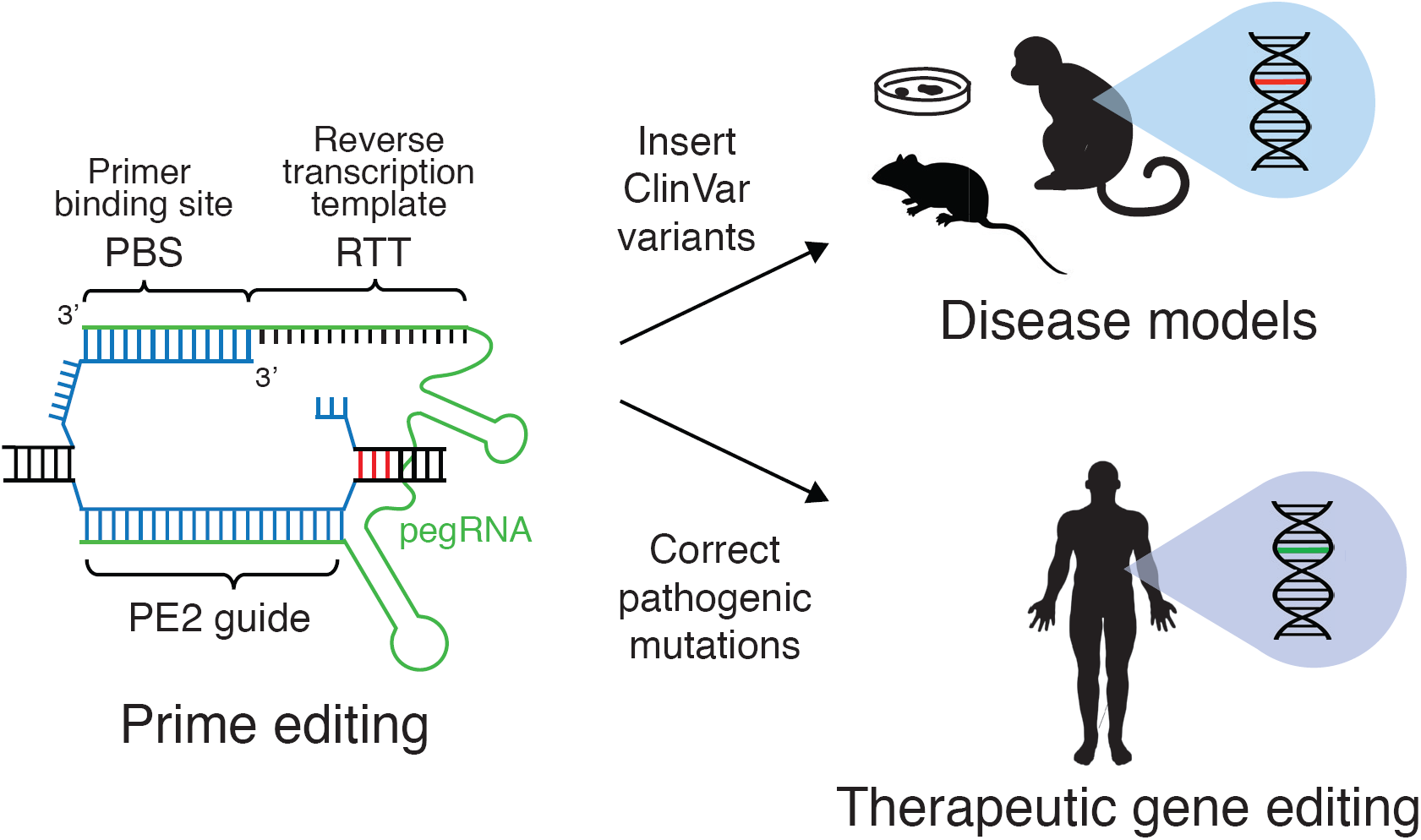
Structure of the prime editing guide RNA (pegRNA) and applications. Prime editing reagents (green) are comprised of a single-guide RNA (PE2 guide RNA), the primer binding site (PBS) and the reverse transcription template (RTT). The PBS and PE2 guide are complementary to the target sequence (blue) and a single-stranded nick is induced three nucleotides from the protospacer adjacent motif (PAM, red). The RTT contains the desired edit to be introduced. PE3/3b single-guide RNA is not shown.

PE design consists of four components: the primary single-guide RNA (sgRNA) that will target a specific site for inducing a nick, the primer binding site (PBS), the reverse transcription template (RTT) that includes the desired edit and a secondary sgRNA to improve editing efficiency. The first three components together form the pegRNA (**Fig. 1**), which when used alone is termed the PE2 approach. To further boost PE efficiency, the PE3 and PE3b approaches utilize a secondary sgRNA, that is distinct from the pegRNA, to either induce a nick 40-90 bp downstream the target or within 3 bp downstream of the target, respectively. Importantly, the PE3b approach requires sgRNA sequence complementarity with the edited strand of DNA sequence. Design consideration specifics for PE3 and PE3b both serve to improve the editing efficiency of prime editing by threefold over PE2, with PE3b greatly reducing indel formation^1^.

Given the number of locus-specific reagents required for prime editing, we developed an optimized pipeline for PE design. To ensure ClinVar variants would be targetable within the default PE parameters (see *Methods*), we only examined single base-pair substitutions and insertions and deletions of 10 base-pairs or less, resulting in 66,580 ClinVar variants (39% transition mutations, 25% transversion mutations, 11% insertions and 24% deletions) (**Fig. 2a**). As an initial consideration, we looked for how many pathogenic variants have a Cas9-targetable site; specifically, Cas9 requires a protospacer-adjacent motif (PAM) at the target site to bind and cut. Of these variants, 78% had PAMs within an appropriate distance for PE2 pegRNAs, 72% for PE3 pegRNAs and 16% for PE3b pegRNAs (**Fig. 2a**). When considering PE2 pegRNA design, the least restrictive of the three classes, most variants had at least 1 or 2 available PAMs (**Fig. 2b**) and we designed PE2, PE3 and PE3b pegRNAs wherever possible to correct pathogenic variants (**Supplementary Tables 1-3**).

**Figure 2.**
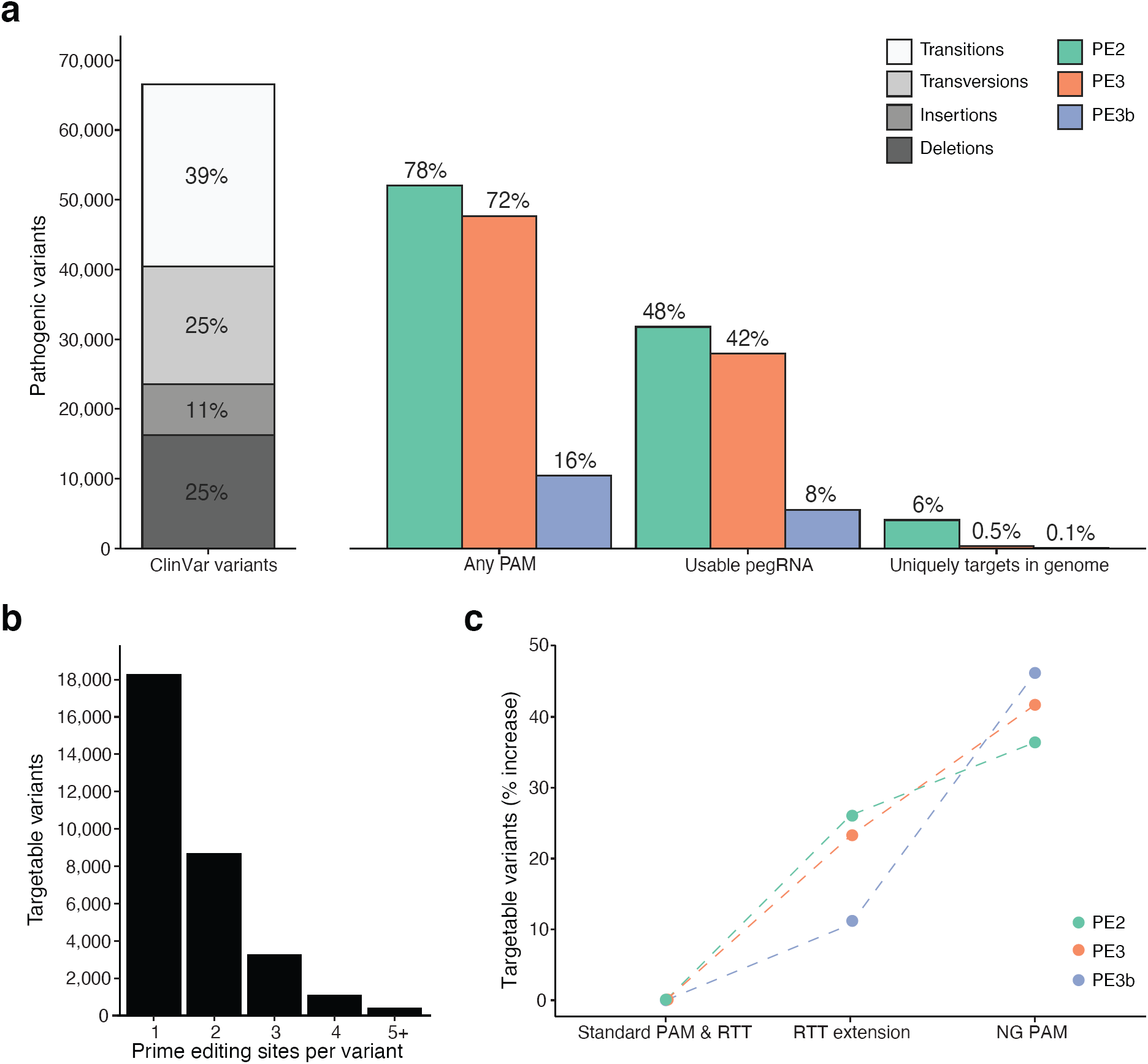
Targetable pathogenic variants with prime editing. **(a)** Total number of ClinVar pathogenic variants by mutation type (transition, transversion, insertion or deletion) *(left)* and the percentage targetable with prime editing after various filtering steps in the design pipeline *(right).* Over 70% of pathogenic variants have protospacer adjacent motifs (PAMs) within a suitable distance for PE2 and PE3 prime editing, however only 16% of pathogenic variants have PAMs within a suitable distance for PE3b prime editing. These percentages decrease as we consider which of the prime editors meet the basic requirements for usability (e.g. no Pol3 terminator motif) and filtering for no predicted off-targets in the genome. **(b)** The number of prime editing sites (PAMs for pegRNA) per pathogenic variant for PE2 prime editors. The majority of pathogenic variants have at least 1 or 2 sites for designing prime editors. **(c)** The increase in the percentage of targetable variants when extending the RTT from 16 nucleotides to 50 nucleotides, and the increase in percentage when allowing for flexible NG PAM recognition instead of *Sp*Cas9 NGG PAM recognition.

Although the PAM site is a requirement, there are several other aspects of pegRNA design, such as suitable GC-content and avoidance of Pol3 terminators motifs, that further constrain the space of targetable sites in the genome. The percentages of targetable variants are reduced to 48%, 42% and 8%, respectively, when only considering usable pegRNAs (**Fig. 2a**). Given the importance of assessing off-target effects when performing genome editing experiments and for therapeutic gene editing, we also analyzed the guide RNAs (both the pegRNAs and PE3/3b sgRNAs) to identify potential off-targets genome-wide. Although use of a Cas9 nickase should minimize off-target genome modification, others have found that nickases can indeed result in indel mutation at a low but detectable frequency^4^. Using a strict threshold for sgRNA-selection based on the Hsu *et al*.^5^ off-target score (≥ 90), we observed that the percentage of targetable variants with PE2, PE3 and PE3b pegRNAs dropped to 6.1%, 0.5% and 0.1%, respectively (**Fig. 2a**).

One method proposed by Anzalone *et al*.^1^ to increase the number of targetable variants is to extend the length of the RTT. Therefore, we re-designed pegRNAs with 50 nucleotide RTTs and, as expected, observed an increase in the number of PAMs for PE2 pegRNAs that were sufficiently close to ClinVar variants **(Supplementary Fig. 1a).** The percentage of targetable variants when considering usable pegRNAs with the RTT extension increased by +26% for PE2, +23% for PE3 and +11% for PE3b (**Fig. 2c**). With stringent off-target filtering, the increases are +11% for PE2, +1% for PE3 and +1% for PE3b (**Supplementary Fig. 1c**).

An alternative strategy is to take advantage of recently engineered Cas9 variants with a less-restrictive PAM sequence, such as Cas9-NG, xCas9, and xCas9-NG^6–9^. These variants require only a NG PAM instead of the NGG PAM for *Sp*Cas9 and have been recently described and optimized in a variety of settings. We re-designed pegRNAs with flexible NG PAM recognition and a standard RTT length. In comparison to the RTT extension, we observed even more targetable variants for PE2 pegRNAs when using NG PAMs (**Supplementary Fig. 1b**). When restricting to targetable variants with usable pegRNAs, we observed the highest increases in targetable variants from all strategies: +36%, +42% and +46% for PE2, PE3 and PE3b, respectively (**Fig. 2c**). Currently, there are no tools to predict off-targets for Cas9 variants with NG PAMs, making it challenging to accurately understand the trade-off with potentially increased off-targets. However, it appears that allowing for flexible PAM recognition overcomes restrictions with designing secondary sgRNAs for PE3 and PE3b prime editors, providing a major benefit for these PE approaches in particular.

Given that there are a large number of genetic variants that are relatively common in the general population (minor allele frequency ≥ 1%), PEs may overlap variants that are not disease-causing but that may disrupt the PAM or their sequence complementarity. Since we designed all pegRNAs reported in this study using a human genome reference (GRCh37), we also assessed whether the sequences of the pegRNAs that require complementarity (the primary sgRNA, the PBS and the secondary sgRNA) or respective PAMs mapped to sites with common human genetic variation using the dbSNP catalog^10^. We found that the majority of usable prime editing reagents (> 95%) did not overlap common genetic variation (**Fig. 3**), suggesting they would not need to be modified further and allow for development of universal therapeutic PEs for most variants.

**Figure 3.**
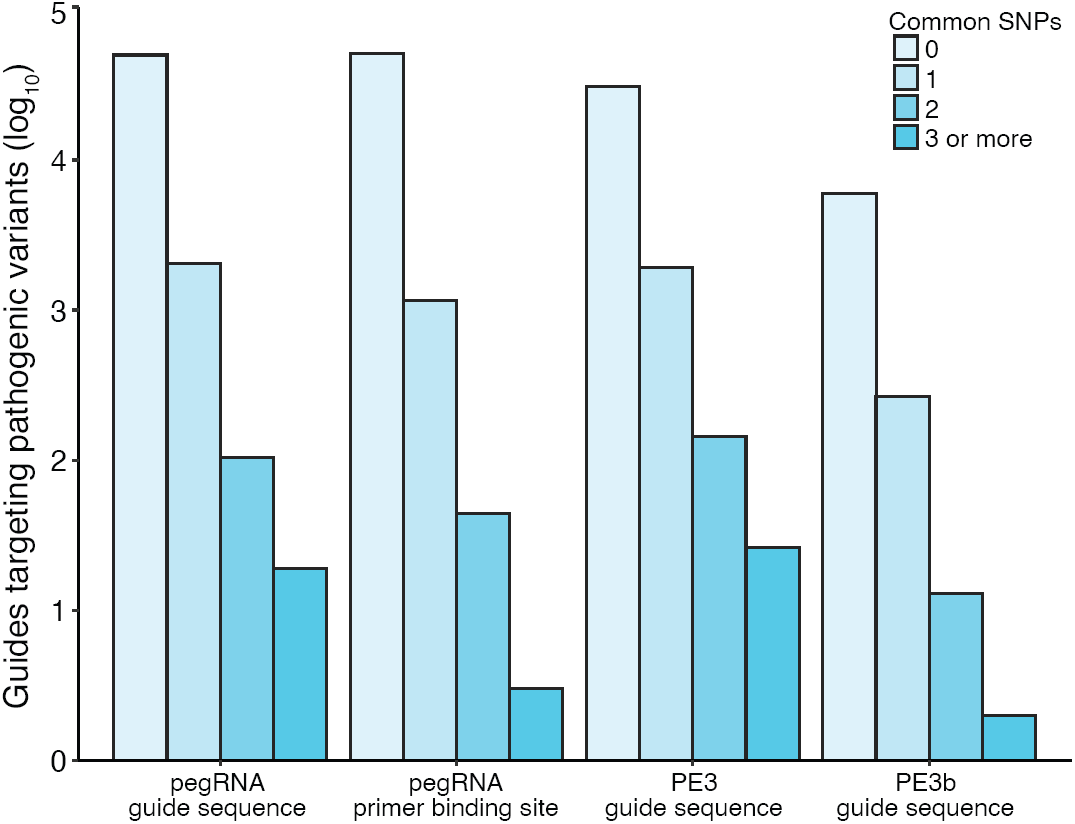
The number of prime editing reagents that map to sites of common human genetic variation. Over 95% of prime editing reagents do not map to sites of common human genetic variation, represented by single nucleotide polymorphisms (SNPs) occurring in over 1% of the gene population. The data are represented on a log scale to allow for visualization of the number of prime editing reagents that do map to sites of common human genetic variation, as these are relatively low in number. However, it should be noted that some prime editors may have to be modified if they map to multiple sites of common human genetic variation.

We have made all designed pegRNAs available through a user-friendly web-based application (http://primeedit.nygenome.org). Using this application, users can search for pegRNAs targeting specific ClinVar variants by their variant ID or by gene ID (if they map to a gene). The pegRNAs discussed in this study were designed to replace ClinVar variants with the human reference allele, however, we also designed pegRNAs to introduce ClinVar variants for creating genetic models of disease (**Supplementary Fig. 2, Supplementary Tables 4-6)**. These alternate pegRNAs for introducing ClinVar variants are also accessible through our web-based application.

Prime editing has tremendous potential for therapeutic gene editing and model organism research. Here, we have shown not only that thousands of known human disease-causing variants are targetable with prime editing reagents, but that thousands more can be targeted with further developments in prime editing and nuclease technology. We have developed an automated pipeline to design prime editing reagents for specific base-pair edits and created a web-based portal to make these reagents readily accessible. Notably, our designed prime editing reagents are for human pathogenic variants from the ClinVar database. Research is still needed to carefully test these prime editing reagents, to determine optimal parameters, but we are now able to design these reagents with ease and at high throughput.

## Acknowledgements

We thank the entire Sanjana laboratory for support and advice. We thank M. Zaran for assistance with the web-tool server. J.A.M. is supported by a Banting Postdoctoral Fellowship and the Canadian Institutes of Health Research. N.E.S. is supported by New York University and New York Genome Center startup funds, National Institutes of Health (NIH)/National Human Genome Research Institute (grant nos. R00HG008171, DP2HG010099), NIH/National Cancer Institute (grant no. R01CA218668), Defense Advanced Research Projects Agency (grant no. D18AP00053), the Sidney Kimmel Foundation, the Melanoma Research Alliance, and the Brain and Behavior Foundation.

## Author contributions

J.A.M and N.E.S. conceived the project and designed the study. J.A.M. wrote the design pipeline and J.R. built the web tool. J.A.M. and J.R. performed analyses. X.G. helped with data presentation. All authors contributed to drafting and reviewing the manuscript, provided feedback and approved the manuscript in its final form.

## Methods

### Automated prime editor design software

We developed a pipeline in R v3.4.4 for designing prime editor guide RNAs (pegRNAs) to induce specific base-pair substitutions, insertions or deletions. This R pipeline uses dplyr v0.8.5, ggplot2 v3.3.0, data.table v1.12.8, BSgenome.Hsapiens.UCSC.hg19 v1.4.0, Biostrings v2.46.0 and future.apply v1.4.0 libraries to design all four components of pegRNAs: the primary sgRNA, the primer binding site (PBS), the reverse transcription template (RTT) and the secondary sgRNA for the PE3 and PE3b classes of prime editors. sgRNA design protocols are well known to design canonical 20 nucleotide (nt) sgRNAs^3^. We reimplemented these sgRNA design considerations to target NGG PAMs. PBS design was recommended by Anzalone *et al*.^1^ to be 13 nt, but can be modified for more favorable GC content of the sequence.

We note here that the PBS end site is explicitly determined by the location of the nick site (3 nt downstream from the PAM). We allowed for the length of the PBS to be determined by user input but set 13 nt as the recommended default. We also provide a measure of the GC content, to be used at the discretion of the user. Anzalone *et al*.^1^ suggested not using PBS sequences if the GC content is outside of approximately 40-60%, therefore we flag any PBS with a GC content less than 35% and greater than 65% as unusable. The default RTT size was set to 16 nt and allows for the targeted edit to be in any position except for the last nt. The PBS and RTT are packaged together and provided as the complete 3’ extension. If secondary sgRNAs are required for PE3 or PE3b classes of prime editors, these are separately generated and can be matched with a corresponding pegRNA complementary to the opposite strand, to ensure nicks are made on opposite strand.

### Identification of human pathogenic variants for prime editing

We identified human pathogenic variants for prime editing with the catalog of variants reported by ClinVar (modified 12/2/2019), available from the NCBI FTP (ftp://ftp.ncbi.nlm.nih.gov/pub/clinvar/vcf_GRCh37/), sorting on the “pathogenic” identifier. We explicitly selected for single base-pair substitutions, insertions less than 10 base-pairs and deletions less than 10 base-pairs, to ensure the full size of any desired edit could be contained with a 16 nt RTT.

### Design of PE2, PE3 and PE3b reagents to target human pathogenic variants

After we identified the subset of human pathogenic variants for prime editing, we generated human reference genome sequences and designed PE2, PE3 and PE3b reagents to introduce the pathogenic allele. We then generated human genome reference sequences with the pathogenic allele instead of the reference allele and designed PE2, PE3 and PE3b reagents to correct these variants. We discuss the reagents used to correct variants throughout this study but provide reagents to introduce the variants as well.

### Identification of prime editing reagents that map to sites of common human genetic variation

One concern when performing genome editing is the occurrence of mismatches between the target genome and the reference from which sgRNAs are designed. To address this, we used dbSNP build 151^10^ to generate a list of all known single nucleotide polymorphisms (SNPs) with minor allele frequencies (MAF) ≥ 1% in the human population. We used bedtools v2.29.2^11^ to perform an intersection between our designed prime editing reagents and common human genetic variation.

### Data availability

The latest release of the ClinVar database is available from the ClinVar FTP (https://ftp.ncbi.nlm.nih.gov/pub/clinvar/). All code and software to reproduce our analyses are available for download from our GitLab repository (https://gitlab.com/sanjanalab/primeediting). All pre-designed prime editing reagents are available for download from a dedicated website (http://primeedit.nygenome.org/). Other data and materials are available from the corresponding author upon reasonable request.

