## Supplementary Figures for "Automated design of CRISPR prime editors for thousands of human pathogenic variants"

**Supplementary tables**

**Supplementary Table 1.** PE2 pegRNAs for correcting ClinVar variants.

**Supplementary Table 2.** PE3 pegRNAs for correcting ClinVar variants.

**Supplementary Table 3.** PE3b pegRNAs for correcting ClinVar variants.

**Supplementary Table 4.** PE2 pegRNAs for introducing ClinVar variants.

**Supplementary Table 5.** PE3 pegRNAs for introducing ClinVar variants.

**Supplementary Table 6.** PE3b pegRNAs for introducing ClinVar variants.

**Supplementary figures**

**Supplementary Figure 1.** Targetable pathogenic variants with prime editing when using a RTT extension or NG PAM.

**Supplementary Figure 2.** Targetable sites to introduce pathogenic variants with prime editors.

### Supplementary Figure 1

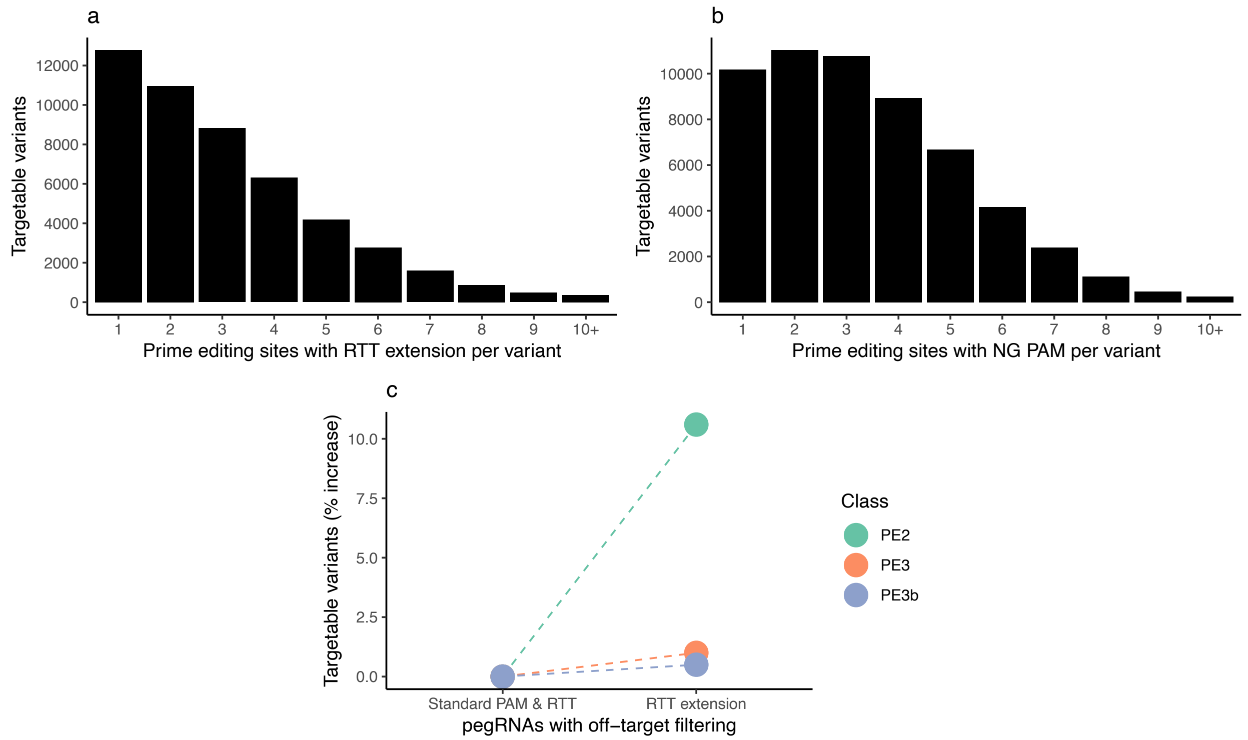


**Supplementary Figure 1. Targetable pathogenic variants with prime editing when using a RTT extension or NG PAM.** The number of prime editing sites (PAMs) per pathogenic variant when considering PE2 prime editors with 50 nucleotide RTTs (**a**) and with flexible NG PAM recognition (**b**). **(c)** The increase in the percentage of targetable variants when extending the RTT length from 16 nucleotides to 50 nucleotides and filtering for no predicted off-targets in the genome.

**Supplementary Figure 2**

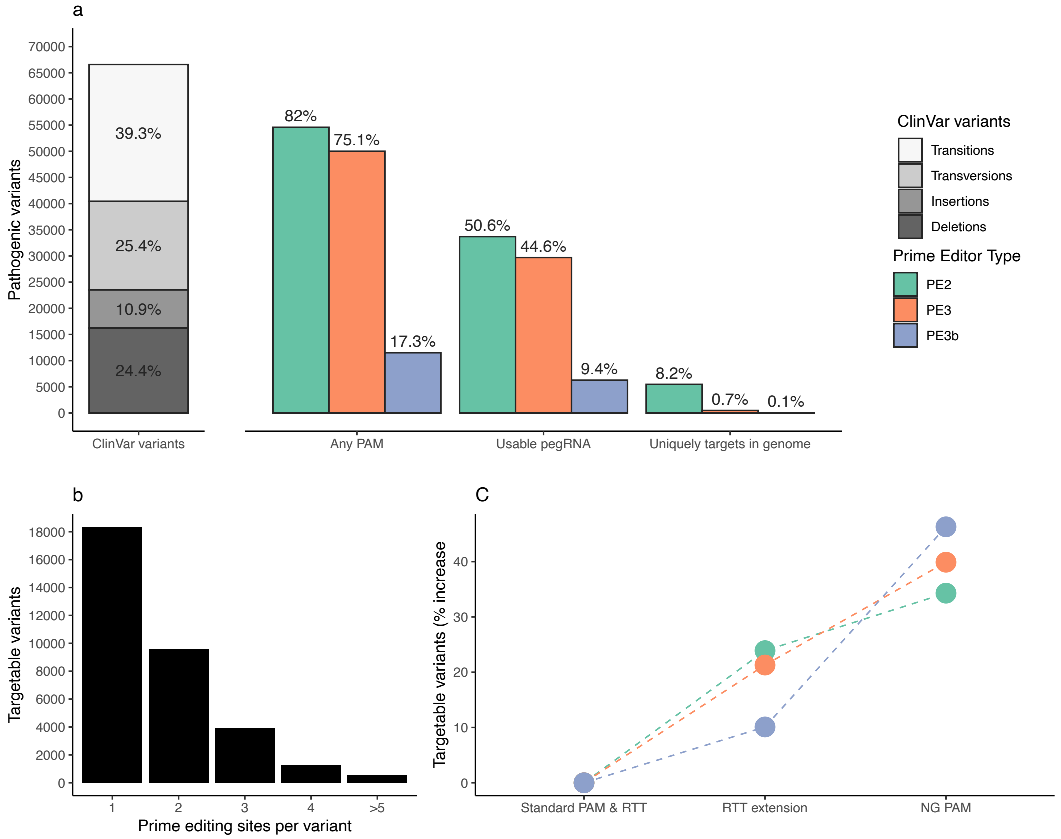


**Supplementary Figure 2. Targetable sites to introduce pathogenic variants with prime editors. (a)** Total number of ClinVar pathogenic variants, broken down by mutation type (transition, transversion, insertion or deletion) and the decrease in the percentage that can be introduced with prime editing under different conditions. Over 70% of target sites have PAMs within a suitable distance for PE2 and PE3 prime editing, however only 16% of target sites have PAMs within a suitable distance for PE3b prime editing. These percentages decrease as we consider which of the prime editors meet the basic requirements for usability (e.g. no Pol3 terminator motif) and filtering for no predicted off-targets in the genome. **(b)** The number of prime editing sites (i.e. PAMs) per target site when considering PE2 prime editors. The majority of target sites for introducing pathogenic variants have at least 1 or 2 sites for designing prime editors. **(c)** The increase in the percentage of targetable sites when extending the RTT from 16 nucleotides to 50 nucleotides, and the increase in percentage when allowing for flexible NG PAM recognition instead of *Sp*Cas9 NGG PAM recognition.
